# Beta 1 Integrin Signaling Mediates Pancreatic Ductal Adenocarcinoma Resistance to MEK Inhibition

**DOI:** 10.1101/862482

**Authors:** Arthur Brannon, Donovan Drouillard, Nina Steele, Shadae Sutherland, Howard C. Crawford, Marina Pasca di Magliano

## Abstract

Pancreatic cancer, one of the deadliest human malignancies, has a dismal 5-year survival rate of 9%. The high mortality rate can be attributed to multiple factors, including late diagnosis and lack of effective therapies. *KRAS* is the most commonly mutated gene in pancreatic cancer, but clinical agents that directly target mutant KRAS are not available. Several effector pathways are activated downstream of oncogenic Kras, including MAPK signaling. MAPK signaling can be inhibited by targeting MEK1/2; unfortunately, this approach has been largely ineffective in pancreatic cancer. Here, we set out to identify mechanisms of MEK inhibitor resistance in pancreatic cancer using primary mouse and human 3D organoid cultures. We optimized the culture of pancreatic tumor organoids that utilized Matrigel as a basement membrane mimetic, facilitating polarized growth. Pancreatic tumor organoids recapitulated mutant KRAS dependency and recalcitrance to MEK inhibition. Treatment of the organoids with trametinib, a MEK inhibitor, had only a modest effect on these cultures. We observed that cells adjacent to the basement membrane mimetic Matrigel survived MEK inhibition, while the cells in the interior layers underwent apoptosis. Our findings suggested that basement membrane attachment provided survival signals. We thus targeted integrin β1, a mediator of extracellular matrix contact, and found that combined MEK and integrin β1 inhibition bypassed trametinib resistance. Our data support exploring integrin signaling inhibition as a component of combination therapy in pancreatic cancer.

## Introduction

Pancreatic Ductal Adenocarcinoma (PDAC), accounting for 90% of pancreatic neoplasms, is projected to become the 2nd leading cause of cancer death in the US by 2030 [1,2]. Almost 95% of PDAC cases express a mutated form of the GTPase KRAS [3]. Activating mutations in *KRAS* (the most prevalent being *KRAS*^*G12D*^), lead to constitutive, aberrant activation of KRAS and subsequent neoplasia [4]. The Mitogen-activated protein kinase (MAPK) pathway is a downstream effector of oncogenic KRAS and its activation promotes cell growth, survival, and proliferation [5]. While KRAS inhibitors are currently not available, the MAPK signaling pathway can be targeted by multiple FDA-approved agents, many of which target the key kinases MEK1/2 [6, 7]. Inhibition of MAPK signaling blocks the onset of carcinogenesis [8], possibly by interfering with the dedifferentiation of acinar cells to duct-like cells that are susceptible to transformation, a process known as acinar-ductal metaplasia (ADM). MEK inhibition has been tested in pancreatic cancer as a single-agent therapy, as well as in combination with Phosphoinositide Kinase-3 (PI3K) pathway inhibition (targeting another downstream effector of KRAS [9, 10]). Unfortunately, these efforts have failed to demonstrate clinical benefit [11].

MEK inhibition using trametinib is tolerated in the PDAC patient population [10]. We set out to understand mechanisms of resistance to trametinib with the goal to identify potential new combination approaches for pancreatic cancer therapy. Since the resistance to trametinib is observed in tumor cells in isolation, we focused here on the cell-autonomous mechanisms of resistance, using a three dimensional (3D) *in vitro* model of PDAC. In this study, we found that cells adjacent to the basement membrane exhibit a survival advantage over cells lacking ECM signaling when administered a MEK inhibitor. Furthermore, KRAS effector signaling is reduced to only ECM-adjacent cells when given an β1 integrin neutralizing antibody. Lastly, dual blockade of both MEK and β1 integrin significantly increased PDAC cell apoptosis compared to singular inhibition of MEK or β1 integrin. These results indicate that β1 integrin plays an important role in mediating PDAC resistance to MEK inhibition.

## Results

### Establishing a 3D culture model of pancreatic cancer

The iKras*;p53* mouse model of pancreatic cancer mimics the progression of the human disease [12]. In this model, oncogenic Kras^G12D^ (Kras*) expression is regulated by a tet-response element, while mutant p53^R172H^ is constitutively expressed in the pancreas, allowing for inducible and reversible expression of Kras* upon administration or removal of doxycycline (DOX), respectively (Fig. 1a). The generation of cell lines from primary tumors formed in iKras*;p53* pancreata was previously described [13]. Subsequently, iKras*;p53* PDAC cells were passaged and maintained in two-dimensional culture in presence of DOX to maintain expression of oncogenic Kras (Fig. 1b).

**Fig. 1.**
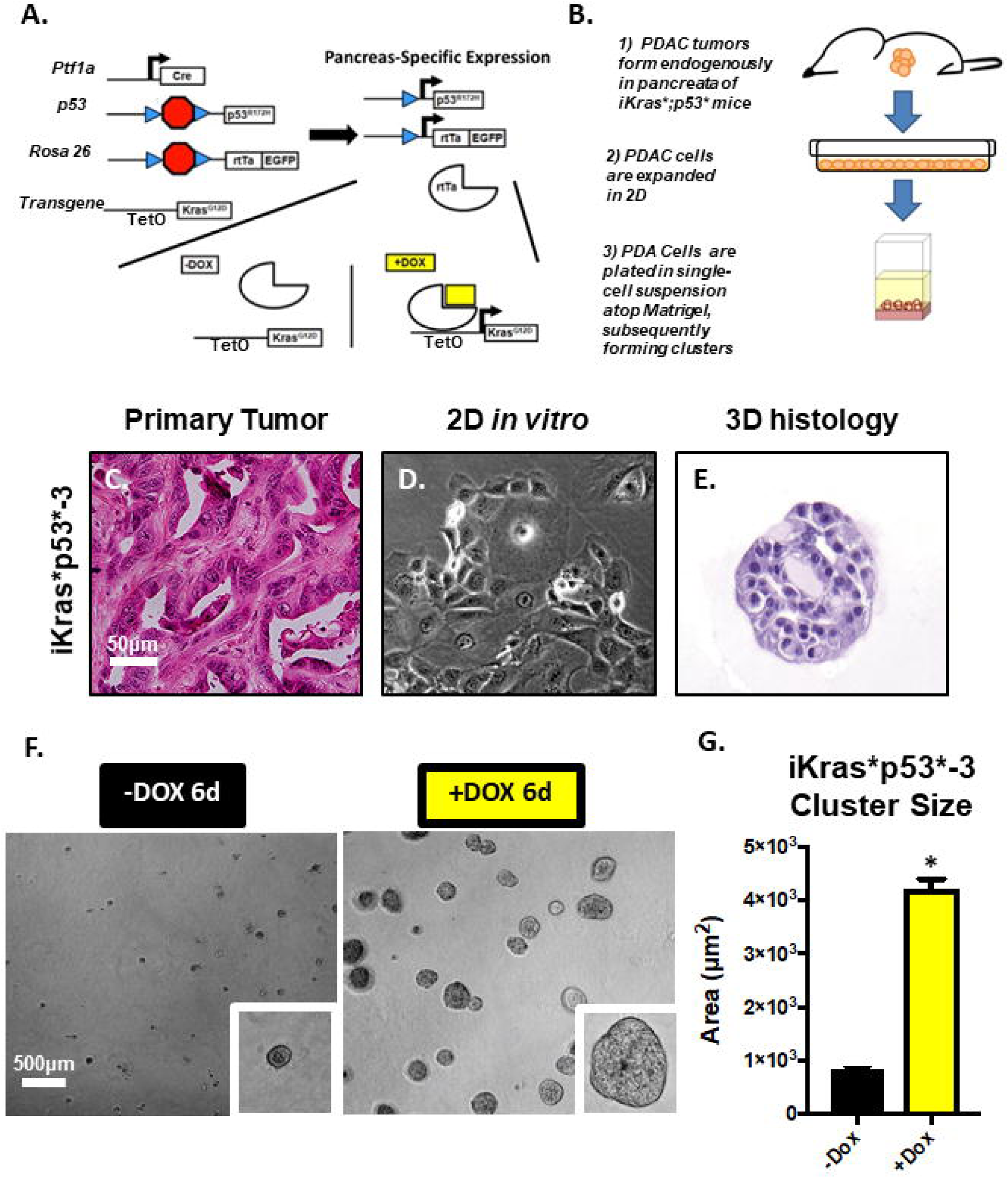
In a 3D culture system, iKras*;p53* cells recapitulate morphologic characteristics of the primary tumor. **a** Schematic describing the genetic model of the iKras*;p53* mouse, wherein administration of doxycycline (DOX) leads to pancreatic-epithelial-cell-specific expression of oncogenic Kras^G12D^ (dominant-negative p53^R172H^ is also constitutively expressed in the pancreatic epithelium). PDA were isolated from endogenous tumors arising. **b** Brief description of endogenous primary tumor formation; In adult mice, DOX was administered through the drinking water. Three days following DOX administration, pancreatitis was induced through two series of intraperitoneal injections of caerulein. Following endogenous tumor formation, tissue was harvested from the primary tumor and the cells were isolated and placed in medium containing DOX. **c** Hematoxylin/eosin stain of primary iKras*p53* PDAC tumors. **d** Brightfield images of PDAC cell lines in 2D culture, maintained in doxycycline (1µg/mL) (Kras* on). **e** Hematoxylin/eosin stain of iKras*p53* PDAC cell cross sections, 6 days following plating in the “on-top” 3D system (cells were also maintained in doxycycline (1µg/mL). **f** Brightfield images of iKras*p53* cells plated in 3D in the absence or presence of doxycycline (1µg/mL) (Kras* on or off, respectively), 6 days following plating of cells. **g** Quantification of cluster area size, 6 days following plating thin the absence (black bars) or presence (yellow bars) of doxycycline (1µg/mL). In quantification, at least 100 clusters were traced and quantified in combined duplicate treatment wells. Bars represent average cluster area ± SD. *p<0.01 in Student’s t test analysis. Scale bars = 50µm; 500µm (low magnification)

Growth in 3D was achieved by trypsinizing and resuspending PDAC cells in single-cell-suspension with cell culture medium that contained DOX and solubilized Growth Factor Reduced (GFR) Matrigel, a basement membrane mimetic comprised of extracellular matrix (ECM) proteins: laminins, Type IV collagen (Col4), and entactin. Matrigel facilitates 3D proliferation and adhesion of cells, interactions that may regulate crucial aspects of cancer molecular pathogenesis *in vivo* [14]. Subsequently, PDAC cells in suspension were plated atop a layer of solidified GFR Matrigel. Approximately 6 days following plating, PDAC cells organized into 3D, discrete clusters that were fixed for histochemical analysis (Fig. 1c-e). To determine whether cells in this 3D assay recapitulated oncogenic Kras* dependency observed in vivo, cells were randomized into two experimental groups: DOX was either withheld or administered to the media at the time of plating to inactivate or activate oncogenic Kras* expression, respectively. Brightfield microscopy was used to monitor cell growth and ImageJ was used to measure PDAC cluster area (Supplementary Fig. 1). DOX administration induced an approximate 4-fold increase in average cluster area (Fig. 1f, g, Supplementary Fig. 2), indicating that oncogenic Kras* expression was sufficient to facilitate tumor cell growth and proliferation in this model. These results are consistent with previous *in vivo* and *in vitro* findings [13].

**Fig. 2.**
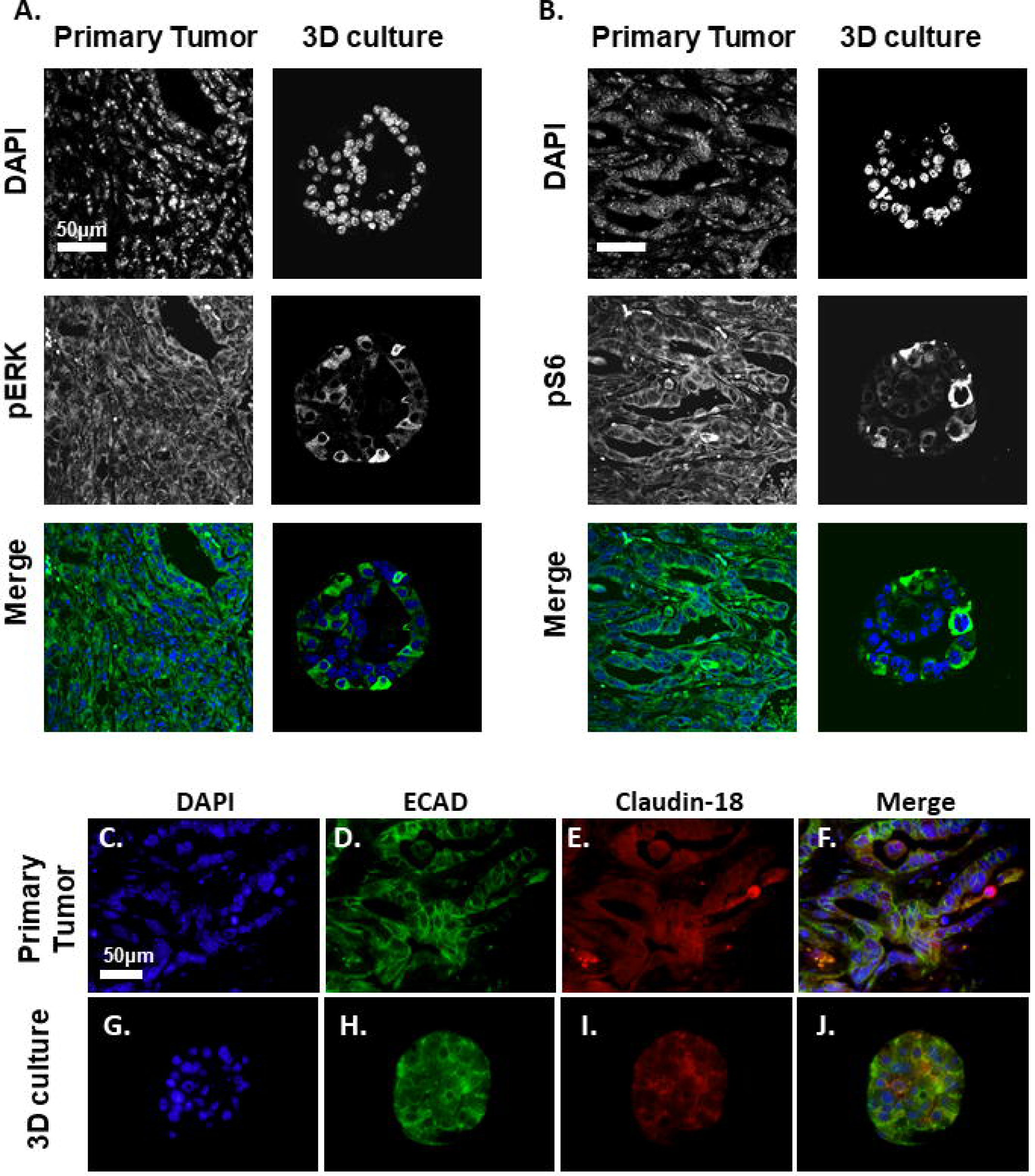
iKras*p53* PDAC cells recapitulate oncogenic Kras* effector pathway activation and expression of membrane proteins in vitro. **a** Nuclear visualization with DAPI along with staining for phosphorylated ERK (pERK) species, indicating MAPK activity in primary tumors and iKras*p53* PDAC cells grown in 3D culture for six days. Scale bar: 50µm. **b** Nuclear visualization along with staining for phosphorylated S6 ribosomal protein (pS6) in primary tumors and PDAC cells in 3D culture. **c-j** DAPI staining along with visualization of membrane proteins e-cadherin and claudin-18 in primary tumor and PDAC cells in 3D culture.

In 2D, oncogenic Kras had a limited effect on cell viability (Supplementary Fig. 3). Conversely, 3D growth was dependent on the expression of oncogenic Kras (Supplementary Fig. 2), mimicking the requirement for oncogenic Kras *in vivo*. We used the 3D system to study the effect of inhibiting Kras* downstream effector pathways, specifically MAPK and PI3K, in the tumor cells. First, we performed immunostaining on iKras*p53* PDAC clusters grown in the presence of DOX for 6 days. Similar to human PDAC and tissue from the primary iKras*p53* tumor (Fig. 2 a, b), iKras*p53* PDAC cells in 3D culture upregulated phosphorylated ERK (pERK) (Fig. 2a) and phosphorylated S6 (pS6) (Fig. 2b), indicating activation of the MAPK and PI3K pathway, respectively. Furthermore, we observed expression of membrane proteins E-Cadherin and Claudin-18 (Fig. 2c-j), both epithelial cell markers. Taken together, these data suggest two implications for our system. First, iKras*p53* PDAC cells in our 3D culture system recapitulate common biological characteristics of human PDAC and other murine *in vivo* models. Second, it can be used to study the effect of signaling pathways in a system that mimics the spatial relationships of tumor cells, while allowing us to dissect signaling, in absence of the complexity of the microenvironment.

**Fig. 3.**
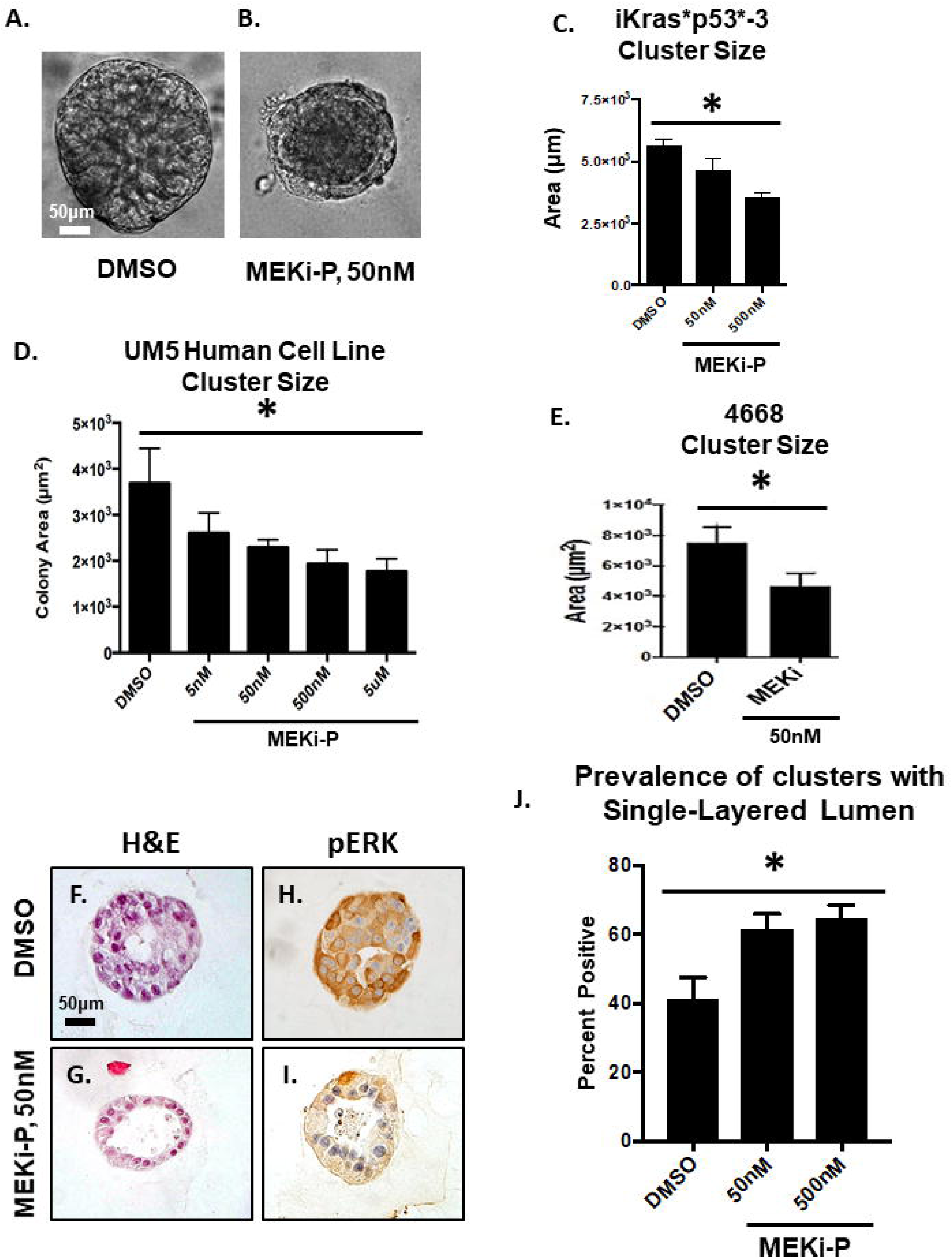
Kras* effector blockade abrogates Kras*-mediated growth in 3D. **a-b** Brightfield images of iKras*p53* cells plated in 3D given either DMSO or a MEK inhibitor for seven days. Quantification of cluster area, 4 days post treatment; at least 100 clusters per treatment group were analyzed. Bars represent cluster area mean ± SEM. ANOVA statistical analysis; * indicates p<0.01 of the cluster area. **c** Quantification of cluster cross sections with a single layer of epithelial cells adjacent to the Matrigel as well as lumen that accounts for >75% of the cluster area. Over 50 cluster cross sections per treatment group were analyzed. Bars represent single layer cluster prevalence per technical replicate, mean ± SEM. ANOVA statistical analysis; * indicates p<0.01. **d, e** Quantification of cluster cross sections of human UM5 cell lines and 4668 cell lines. **f, g** Hematoxylin/eosin stains of representative cluster cross sections of of iKras*p53* cells plated in 3D given either DMSO or a MEK inhibitor for seven days. **h, i** Immunohistochemical staining of phosphorylated ERK at Thr202 and Tyr204 (pERK); brown dye indicates positive staining. Scale bars: 50µm.

### MEK inhibition inhibits PDAC growth in 3D and induces apoptotic lumen formation

To inhibit MAPK signaling, clusters were grown for 6 days and then administered PD325901 (MEK inhibitor, abbreviated MEKi-P) for 4 days; DOX was present in the media at all points, so that oncogenic Kras* was constitutively expressed. Administration of the inhibitor was sufficient to abrogate Kras*-mediated cluster growth (Fig. 3a-e). Upon histologic analysis we found that MEK inhibition decreased the expression of pERK, suggesting successful inactivation of the MAPK signaling pathway (Fig. 3f-i). However, similar to the in vivo situation, cultures survived, indicating resistance to MEK inhibition. The MEK-resistant cells formed single-layer clusters, while the lumen (occupying >75% of cross-section area) contained apoptotic debris. This finding was dose-dependent (Fig. 3j), as increasing concentration of MEKi led to increased prevalence of single-layered clusters with large lumens. These results suggested that cells at the periphery of the cluster, and thus in contact with the basement membrane, were uniquely resistant to MEK inhibition. Intriguingly, this finding recapitulated what we and others observed in vivo upon inactivation of oncogenic Kras [13, 15].

To confirm this initial finding with a different MEK inhibitor, PDAC cells were grown for 1 week and administered a clinically available MEK inhibitor, trametinib (MEKi-T) for 4 days. To visualize any morphologic changes induced by MEK inhibition, PDAC clusters were fixed, sectioned, and stained for hematoxylin and eosin. Similar to clusters treated with MEKi-P, MEKi-T treated clusters were found to have an apoptotic lumen--evidenced by unorganized hyaline aggregation as well hyperchromatic debris, which indicates nuclear fragmentation, features consistent with cells that undergo programmed cell death (Fig. 4c). The prevalence of clusters with apoptotic debris that occupied >75% of a single lumen was found to be significantly increased approximately 7-fold following MEK-T administration (Fig. 4d). To establish that the cells had indeed undergone apoptosis, we stained sections of PDAC clusters for cleaved caspase-3 (CC3) (Fig. 4a, b). Thus, using two different MEK inhibitors, we showed that cancer cells vary in their sensitivity to MEK inhibition.

**Fig. 4.**
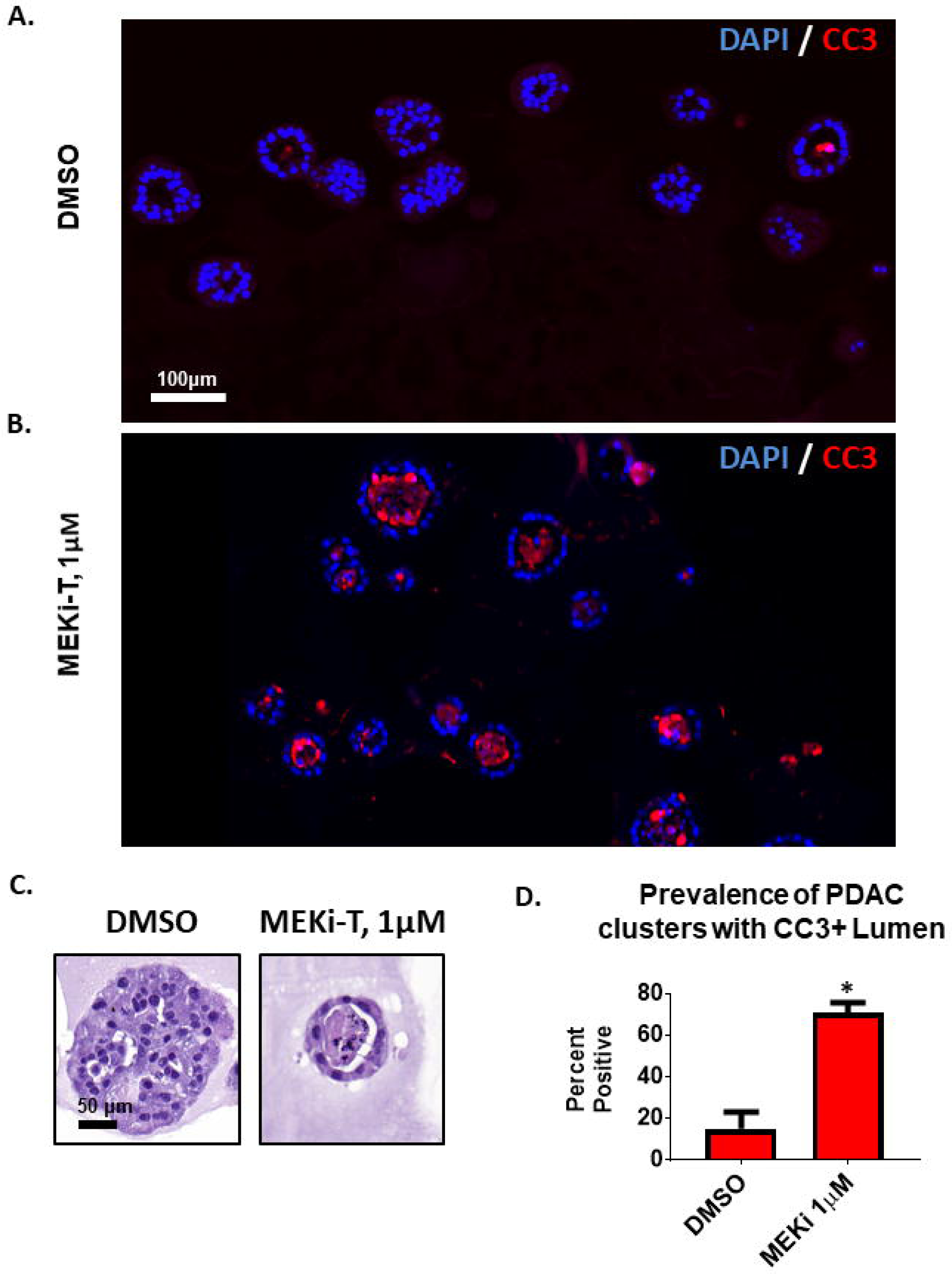
MEK inhibition induces apoptotic lumen formation. iKras*p53* PDAC cells (9805) were grown in 3D culture for 6 days and then treated with the MEK inhibitor trametinib (MEK-T) for 4 days (Kras*on the entire experiment). **a, b** Immunofluorescence staining of cleaved caspase 3 (CC3) in red, indicating apoptosis. Scale bar: 100µm. **c** Hematoxylin/eosin staining of representative PDAC cluster cross sections from corresponding treatment groups. Scale bar: 50µm. **d** Quantification of prevalence of PDAC cluster cross sections wherein the lumen accounts for >75% of the cluster and contains CC3-positive debris. Bars represent mean prevalence ±SEM of over 50 cluster cross sections in 3 grouped biological repeat studies. Statistics: student’s t test; * indicates p<0.001.

### PDAC cells adjacent to Matrigel display a survival advantage

The prevalence of single-layer cluster morphology suggests that cells in contact with the Matrigel basement membrane mimetic have a survival advantage. In this 3D system, Matrigel coats the entire outside of the cluster *in vitro*, so following fixation and sectioning, the outermost layer of cells in cluster cross-sections are considered adjacent to the Matrigel (Supplementary Fig. 1). This can be visualized by staining for Type IV collagen (Col4), a primary component of Matrigel and the basement membrane *in vivo*. Staining reveals Col4 enrichment on the outer edge of clusters treated with either vehicle (DMSO) or MEKi-T, suggesting that MEK inhibition does not affect organization of basement membrane (Fig. 5a). Upon quantification of cells adjacent or nonadjacent to Matrigel, the MEK-T treated clusters showed an increased prevalence of cells adjacent to the Matrigel, suggesting that these cells have a survival advantage over cells that lack ECM attachment (Fig. 5b). The phenomenon of anoikis, a form of programmed cell death in response to loss of ECM signaling, is well described in other systems. [16]. We hypothesized that Matrigel-adjacent cells were able to resist anoikis due to their ability to interact and exchange signals with ECM components.

**Fig. 5.**
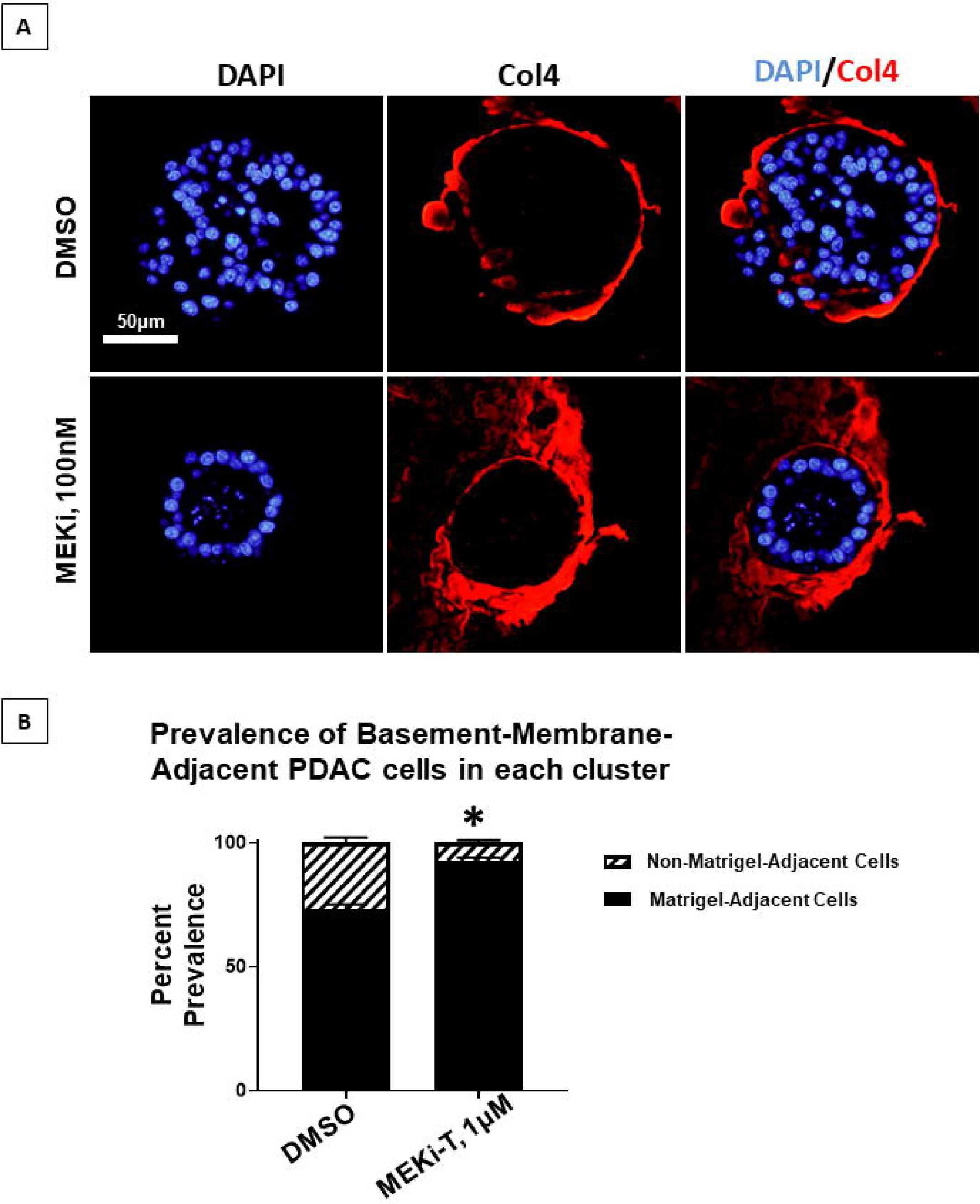
Matrigel-adjacent PDA cells display a survival advantage. iKras*p53* PDAC cells (9805) were grown in 3D culture for 6 days and then treated with the MEK inhibitor trametinib (MEK-T) for 4 days (Kras*on the entire experiment). **a** Representative PDAC cluster cross sections, immunofluorescence staining of type IV collagen (Col4) in red. Scale bar: 50µm. **b** Quantification of the prevalence of non-matrigel-adjacent versus matrigel-adjacent, DAPI-positive nuclei in striped and black bars, respectively. Bars represent mean prevalence ±SEM of over 50 cluster cross sections in 3 grouped biological replicate studies. Statistics: student’s t test; * indicates p<0.001.

Human and murine PDAC cells have been shown to engage in complex signaling with the surrounding microenvironment, yet the implications of this signaling and their effects on disease pathogenesis are currently poorly understood. In multiple systems, the transmembrane, bidirectional signaling molecule β1 integrin has been implicated in coordinating cell-to-cell and cell-to-ECM interactions [17]. And in the normal pancreas, β1 integrin expression is necessary for acinar maintenance [18]. In our 3D system, we found that iKras*p53* PDAC cells expressed β1 integrin (Supplementary Fig. 4), and this expression was not affected by administration of MEKi-T (Supplementary Fig. 4). Given that Matrigel-adjacent PDAC cells show a survival advantage and express β1 integrin, we hypothesized that β1 integrin signaling mediated survival in this population.

### Beta-1 integrin inhibition disrupted cell:cell organization and decreased Kras effector signaling

The function of β1 integrin is highly dependent on cell type as well as the cell’s immediate microenvironment [19, 20]. To determine the functional importance of β1 integrin signaling in our 3D culture model, PDAC clusters were grown for 7 days and treated with either solubilized rat immunoglobulin G (IgG) or a β1 integrin neutralizing antibody [Supplementary Table 1] for 4 days. Because antibodies are targeted to an extracellular epitope, and are unable to cross the cell membrane, the β1 integrin neutralizing antibody specifically blocks β1 integrin outside-in signaling. Following administration of either IgG or β1 integrin neutralizing antibody, cells were fixed, sectioned histologically, and analyzed.

After 4 days of treatment with control IgG, brightfield microscopy of PDAC clusters showed distinct colonies with smooth and ordered interactions with the surrounding Matrigel (Fig. 6a). In contrast, β1 integrin blockade induced structural disorder of PDAC clusters, disrupting their interaction with surrounding Matrigel as well as induced some disintegration of clusters, as evidenced by an increase in scattered, single cells (Fig. 6b). Upon histological analysis, PDAC clusters treated with IgG formed organized colonies with 1 or multiple lumen (Fig. 6 c-f); conversely, β1 integrin blockade abrogated lumen formation (Fig. 6 g-j), suggesting that β1 integrin outside-in signaling is necessary for PDAC cell:cell adhesion and formation of higher ordered 3D structure. Moreover, β1 integrin blockade affected E-cadherin (ECAD) expression at the cell membrane. PDAC clusters in the treated group showed punctate, noncontiguous staining, indicating disruption of ECAD localization to cell membranes (Fig 6d-f, h-j).

**Fig. 6.**
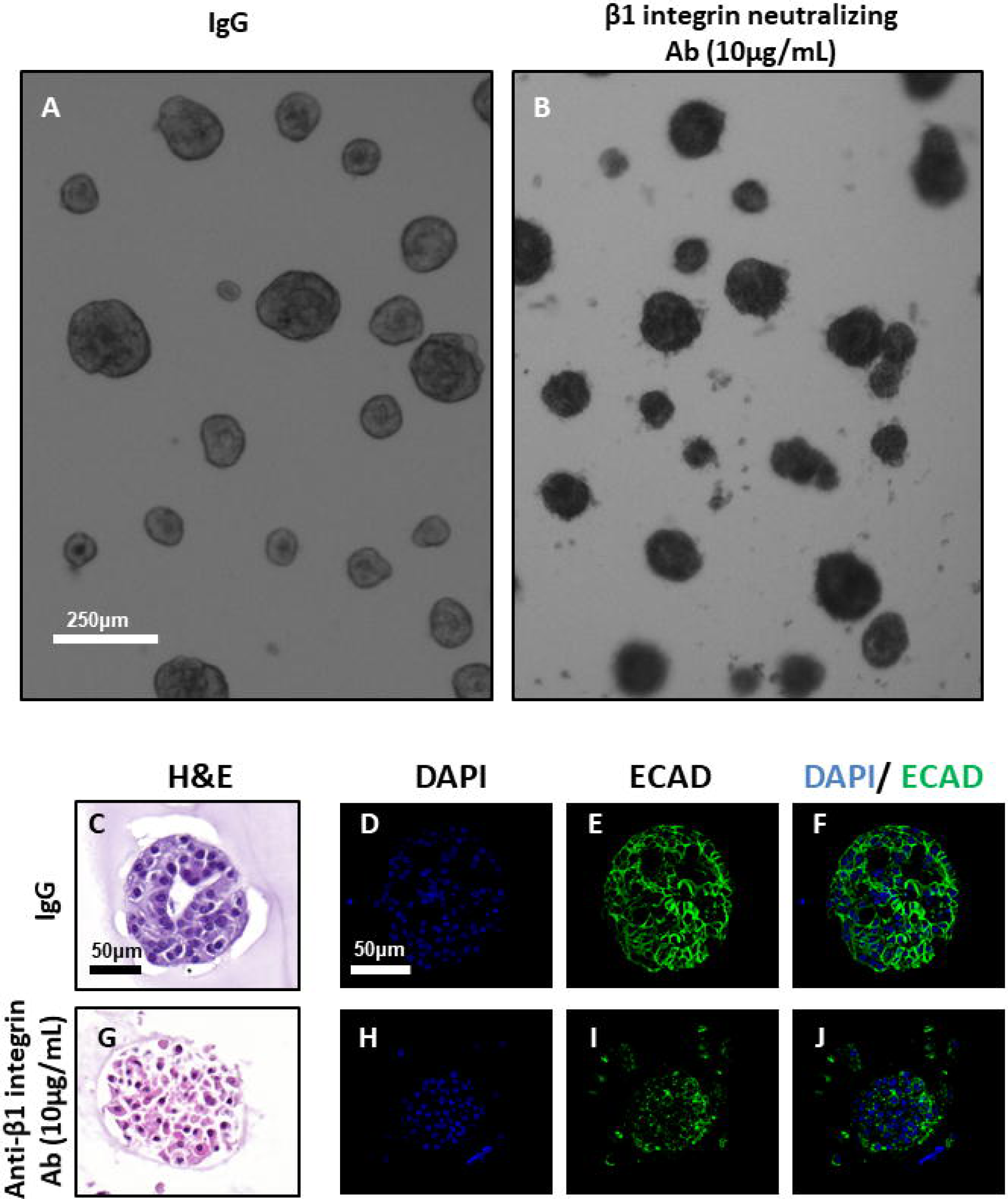
Beta 1 integrin blockade disrupts membrane dynamics. iKras*p53* 9805 PDAC cells were grown in 3D for 7 days and subsequently treated for 4 days with either vehicle IgG or anti-β1 neutralizing antibody (10µg/mL). **a, b** Brightfield, low-magnification microscopy of 9805 PDAC clusters in vitro. Scale bar: 250µm **c, g** Hematoxylin/eosin staining of PDAC clusters cross sections, treated with either control (IgG) or anti-β1 integrin blocking antibody Scale bar: 50µm. **d-f, h-j** Immunofluorescence of PDAC cluster cross sections. Scale bar: 50µm. **d, h** Nuclear visualization with DAPI; **e, i** Stain indicating e-cadherin (ECAD) localization; **f, j** Overlay of DAPI and ECAD channels.

We then evaluated the effects of β1 integrin blockade on Kras* effector signaling. Following administration of control IgG, PDAC clusters demonstrated activation of MAPK and PI3K signaling in Matrigel-adjacent and non-Matrigel-adjacent cells indicated by positive staining of pERK and pS6 (Fig. 7 b, l, Supplementary fig. 4). Strikingly, following β1 integrin blockade, Kras* effector signaling was restricted to Matrigel-adjacent cells (Fig. 7 g, q). Though β1-integrin-blocked PDAC clusters showed some localization of β1 integrin to the ECM (Fig. 7 h, r), colocalization of β1 integrin and pERK or pS6 signals was rare (Fig. 7 i, s). These results suggest that β1 integrin signaling is necessary for PDAC upregulation of Kras* downstream signaling in the absence of ECM signaling in our system. Furthermore, β1 integrin outside-in signaling is dispensable for Kras* effector signaling, if cells are physically adjacent to the ECM.

**Fig. 7.**
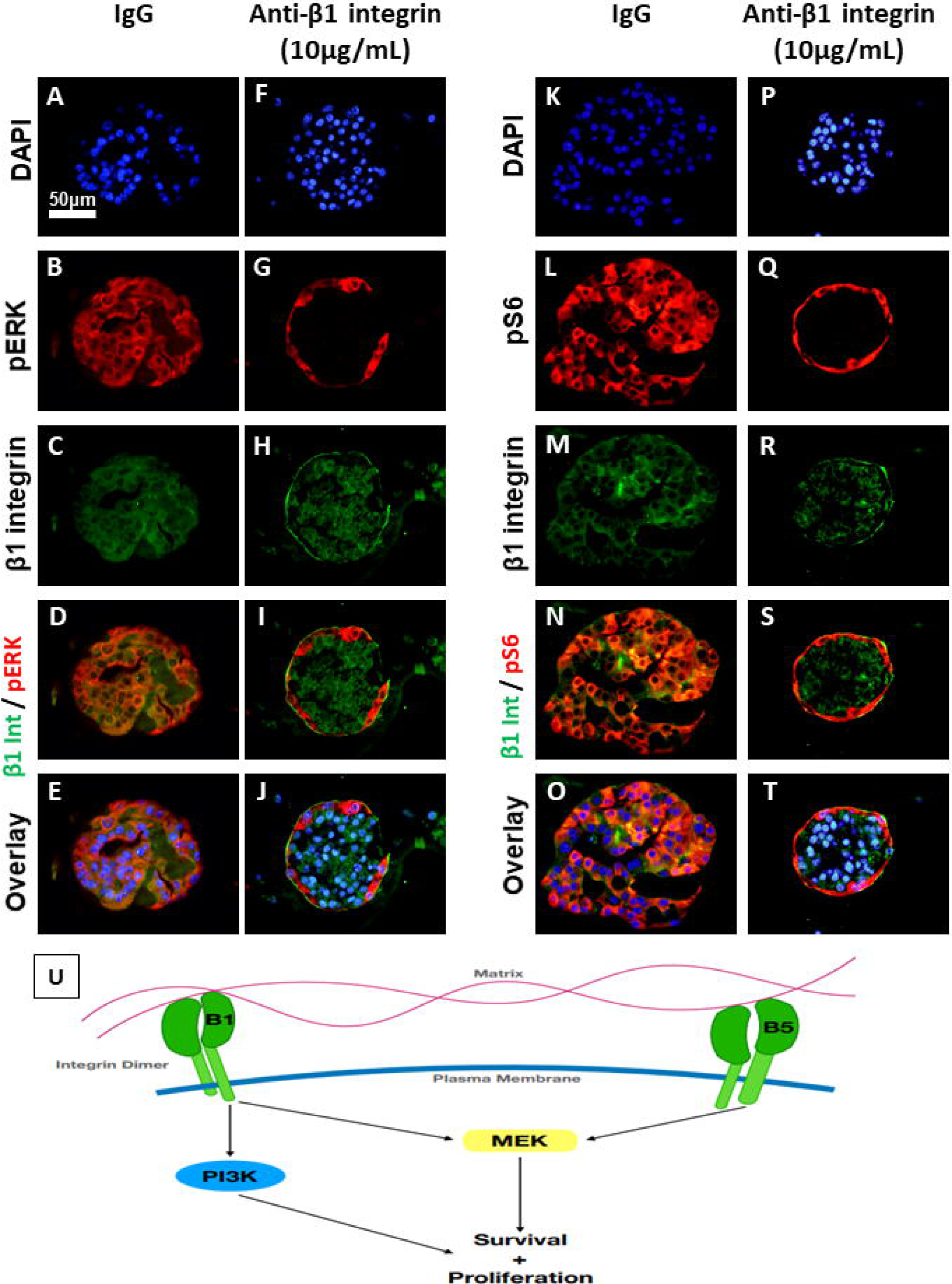
β1 integrin blockade decreases Kras* effector signaling and restricts activation to Matrigel-adjacent cells. iKras*p53* 9805 PDAC cells were grown in 3D for 7 days and subsequently treated for 4 days with either vehicle IgG or anti-β1 neutralizing antibody (10µg/mL). **a-e, k-o** Pictured are representative PDAC cluster cross sections of control treated or anti-β1 integrin-treated cells. Scale bar: 50µm. **b, g** Clusters were stained for phosphorylated ERK (pERK) species, indicating MAPK activity. **l, q** Staining for phosphorylated S6 ribosomal protein (pS6). **c, h, m, r** Staining for β1 integrin. **d, i, n, s** Overlay of red and green channels. **e, j, o, t** Overlay of DAPI, red, and green channels. **u** Schematic outlining the potential signaling pathways involved in β1 and β5 integrin signaling that are responsible for PDAC cell survival and proliferation.

### Dual blockade of MEK and β1 integrin synergize to induce cancer cell death

Prior studies have demonstrated that epithelial cell lumen formation is an active process that requires both cell:cell adhesion and cell:ECM signaling. Since lumen formation appears to be protective for PDAC cells in the context of MEK inhibition (Fig. 4d, 5b) and β1 integrin was necessary for lumen formation in our model (Fig. 6g-j), we hypothesized β1 integrin signaling blockade would prevent lumen formation and potentiate the ability of MEK inhibition to induce apoptosis. To test this, PDAC cells were grown in 3D for 7 days and treated for 4 days with vehicle (DMSO+IgG), MEK-T, an anti-β1 integrin neutralizing antibody, or a combination of the compounds. Subsequently, PDAC clusters were fixed, sectioned and prepared for immunofluorescence to examine CC3 or TUNEL staining. Singular administration of either MEK-T or anti-β1 integrin neutralizing antibody increased apoptosis, as indicated by increased CC3 staining, compared to the vehicle group (Fig. 8a-p, Supplementary Fig. 6). Furthermore, dual inhibition led to significantly increased apoptosis when compared to singular blockade of either pathway (Fig. 8q). TUNEL staining demonstrated similar findings: singular blockade of either MEK or β1 integrin significantly increased cell death, while clusters in the dual blockade group showed significantly increased death when compared to singular blockade of either pathway (Figure 8r, s). These results, taken together, suggest that β1 integrin signaling mediates PDAC resistance to MEK inhibition.

**Fig. 8.**
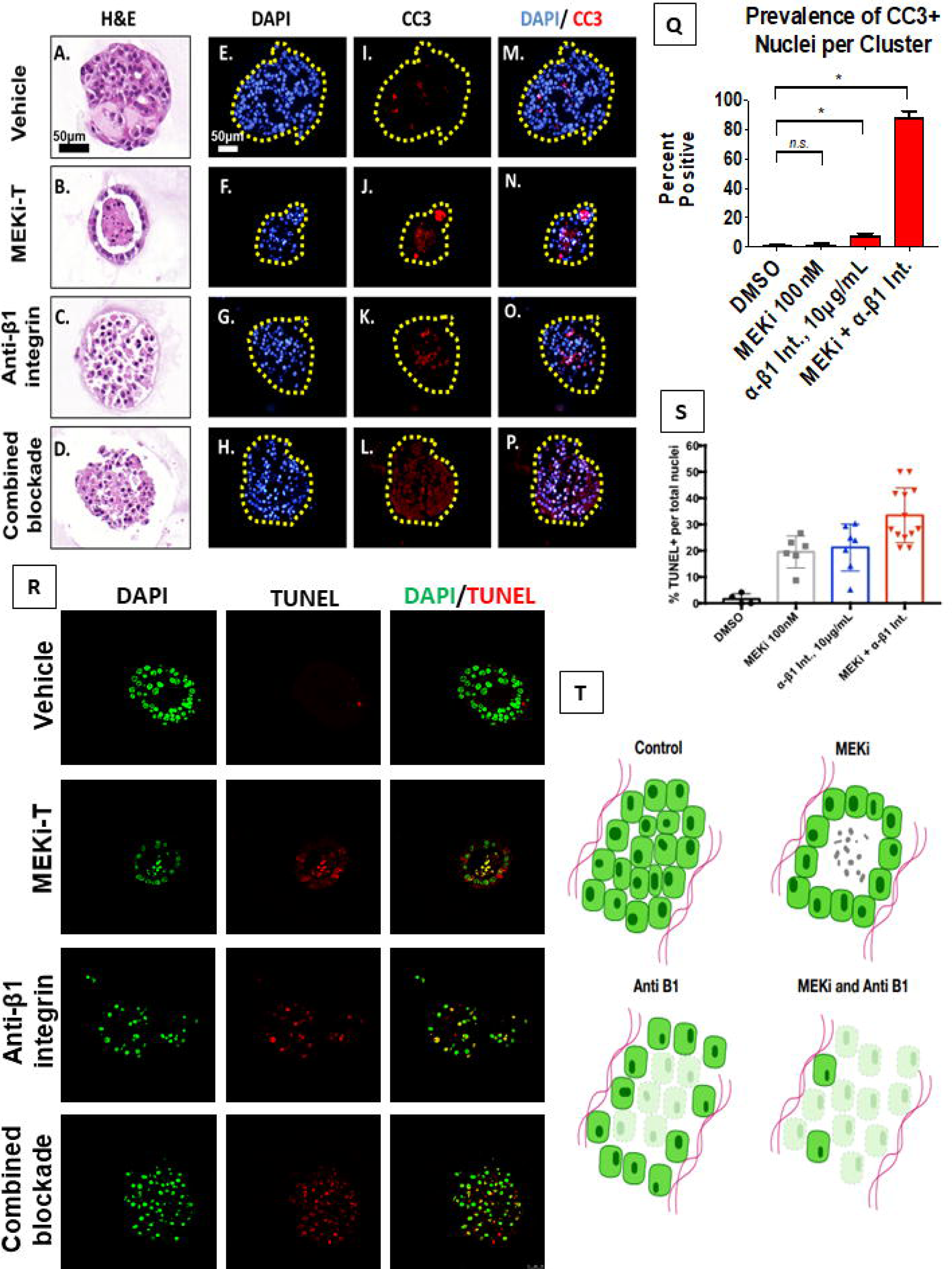
β1 integrin blockade sensitizes PDA cells to apoptosis. iKras*p53* 9805 PDAC cells were grown in 3D for 7 days and subsequently treated for 4 days with vehicle DMSO, trametinib, an anti-β1 neutralizing antibody (10µg/mL), or dual trametinib and anti-β1 neutralizing antibody. **a-d** Hematoxylin/eosin stains of representative cluster cross sections. Scale bar: 50µm. **e-h** Nuclear visualization with DAPI. **i-l** Immunofluorescence staining of cleaved caspase 3 (CC3) in red, indicating apoptosis. **m-p** Overlay of DAPI and CC3 channels. **q** Quantification of CC3-positive nuclei. Bars represent mean prevalence of CC3 positivity per defined nucleus ± SEM. At least 50 unique cluster cross sections from 3 biological replicate experiments were analyzed. Statistical analysis: student’s t test. n.s.=p>0.05; * indicates p<0.001. **r** Single channel and merged channel images of TUNEL immunofluorescent staining of representative cluster cross sections receiving vehicle, single, or dual treatment. **s** Quantification of TUNEL-positive nuclei. Every dot represents a cluster indicating the percentage of TUNEL positive nuclei within the cluster, whereas the horizontal lines are the median percentages of TUNEL positive nuclei throughout multiple clusters. Statistical analysis: One-way anova with post-hoc analysis. * indicates p<0.005, ** indicates p<0.0005. **t** Diagram depicting cluster phenotype upon response to treatment in 3D cell culture. Clusters receiving no treatment will form solid, spheroids. With trametinib administration, cells in the middle of the spheroid will undergo apoptosis leaving a hollow 3D cluster with the remaining living cells attached to the ECM. Clusters receiving the anti-β1 neutralizing antibody will form disorganized clusters with similar preference for ECM attachment. There is an increase in apoptosis in these clusters compared to the control. Clusters receiving dual treatment of trametinib and the anti-β1 neutralizing antibody will again form disorganized clusters with a high percentage of the cells undergoing apoptosis.

## Discussion

Our results show that MEK inhibition induced lumen formation in iKras*p53*-mouse-derived PDAC cells *in vitro*. Furthermore, the presence of Cleaved Caspase 3 and TUNEL positive cells in the lumen suggested that MEK inhibition sensitized PDAC cells to anoikis, programmed cell death in response to detachment from the ECM [24]. Cell:cell adhesion and ECM:cell are key signaling networks that may regulate PDAC progression [21-23]. Elucidation of these pathways may provide insight into mechanisms that regulate disease persistence, which would ideally lead to practical, effective molecular targeting and reduction of PDAC burden. Given that ECM-adjacent PDAC cells displayed a survival advantage in the context of MAPK inhibition, we sought to characterize the role of ECM:cell signaling in our system. We hypothesized that β1 integrin played an important role in the survival of ECM-adjacent PDAC cells.

The integrin family of cell adhesion receptors mediates a multitude of cellular functions crucial to the initiation, progression and metastasis of solid tumors [25]. In human breast cancer, overexpression of β1 integrin is correlated with poor prognosis [26, 27], and in vitro β1 blockade induced apoptosis of malignant cells [28]. In pancreatic cancer, overexpression of β1 is correlated with poorer disease-free survival [29]. In a murine model of pancreatic β cell cancer, genetic ablation of β1 integrin led to reduced tumor cell proliferation and senescence [30]. Pancreatic cancer cell lines overexpress integrin a subunits 1-6 and β subunits; moreover, the β1 subunit is found to be constitutively active, mediating adhesion and invasiveness in some PDAC lines [31]. In our study, we found that lumen formation as well as PDAC cell survival in the context of MEK inhibition were significantly decreased following β1 integrin neutralizing antibody administration, suggesting that PDAC resistance to MEK inhibition is mediated in part by β1 integrin signaling.

Strikingly, singular blockade of β1 integrin decreased Kras* effector signaling in cells lacking ECM attachment but failed to decrease MAPK or PI3K signaling pathway activation in Matrigel-adjacent PDAC cells. These data suggest two novel findings in relation to iKras*p53* PDAC cell biology in vitro. First, cells lacking ECM interaction require β1 integrin to upregulate downstream effectors of Kras*. This is consistent with previous findings that β1 integrin is involved in ERK signaling in PDAC [33] and inhibition promotes central necrosis [34]. The combination therapy likely exacerbated apoptosis due to β1 integrin’s ability to activate PI3K signaling in PDAC [35]. Furthermore, β1 integrin has be shown to be necessary to activate PI3K when MEK is inhibited [36] and PI3K inhibition produces central necrosis in a pattern similar to our data [37]. The second conclusion drawn from our data suggest that ECM-adjacent cells upregulate Kras*-effector signaling in a β1 integrin-signaling-independent manner. One likely mechanism to explain this is β5 integrin signaling (Figure 7u). β5 integrin is present in PDAC [35] and can activate MEK signaling in breast cancer [38]. Upon β5 integrin inhibition in breast cancer, proliferation is decreased [38]. Lastly, suppression of a specific heterodimer containing β5 integrin (α5β5) in colon cancer increases the function of a heterodimer containing β1 [34]. This crosstalk between β1 and β5 integrins suggest that β5 integrin could upregulate Kras-effector signaling in ECM-adjacent cells when β1 integrin is inhibited. However, future tests will need to be done to examine the role of β5 integrin in PDAC as well characterizing how subunit-specific signaling may modulate different characteristics of PDAC interaction with the microenvironment, subsequently regulating PDAC cell survival.

## Methods

### Murine PDAC model and establishment of primary cell cultures

PDAC tumor cell lines from iKras*p53* mice were established as previously described [12]. p48-Cre (Ptf1a-Cre) mice were crossed with TetO-KrasG12D, Rosa26rtTa/rtTa and p53R172H/+ mice to generate p48Cre [38]; TetO-KrasG12D [38, 39]; Rosa26rtTa/+ [38-40]; p53R172H/+ [38-41] (iKras*p53*) mice. In adult mice, DOX was administered through the drinking water, at a concentration of 0.2g/L in a solution of 5% sucrose, and replaced every 3–4 days. Three days following DOX administration, pancreatitis was induced through two series of eight hourly intraperitoneal injections of caerulein (Sigma C9026), at a concentration of 75 μg/kg, over a 48-hour period, as previously described. Following endogenous tumor formation, tissue was harvested from the primary tumor, minced, and digested with 1 mg/ml collagenase V (Sigma) at 37°C for 15 minutes. RPMI-1640 (Gibco) +10% Fetal Bovine Serum +1% penicillin/streptomycin was used to halt digestion and cells were isolated by filtration through a 100 um cell strainer and plated in complete medium containing DOX (Sigma) at 1 μg/mL [40].

### Cell culture

iKras*p53* PDAC cells were maintained and passaged in 2D culture in IMDM media +10%FBS +DOX 1μg/mL. The 3D assay used was adapted from multiple models [14, 42, 43]. Four-well chamber slides (Corning 354104) were coated with 50-100μL Growth-Factor-Reduced (GFR) Matrigel. Afterward, the slides were incubated for 10 minutes at 37°C to induce solidification of GFR Matrigel. PDAC cells were trypsinized and centrifuged. Pellets were resuspended in single-cell-suspension in Waymouth’s media (Gibco 11220-035) +10% Fetal Bovine Serum (Gibco 10438-036) + 1% penicillin/streptomycin (Gibco 15140-122) and the solution was added atop the GFR Matrigel surface. Every 2-3 days, media was changed. In experiments using small molecule inhibitors, inhibitors were solubilized in DMSO or 1x PBS, and vehicle or drug treatment groups were administered in either technical duplicate or triplicate. Representative experiments of at least 3 biological replicates has been shown unless otherwise noted.

### Fixation and staining

PDAC clusters were fixed in formalin at 22°C for at least 2 hours. After fixation, chambers from chamber slides were removed, and PDAC clusters in GFR Matrigel were collected and placed in a cryomold (Sakura 4557) with Histogel (Thermo Scientific HG-4000-012) [44]. These samples were then processed, embedded in paraffin, and cut into 5*μ*m-thick sections. The primary antibodies were added and then antigen retrieval was accomplished using the respective antibodies and concentrations seen in Supplementary Table 1. Histology and immunofluorescence analysis were performed as described below.

### Brightfield and Cleaved-Caspase 3 Imaging and Quantification

Brightfield and fluorescent images were acquired with an Olympus BX-51 microscope DP71 digital camera/software as well as Pannoramic SCAN II slide scanner and software. Brightfield, low-magnification images of clusters growing in chamber slides were used to quantify PDAC cluster area. Pictures of at least 5 non-overlapping areas were taken and a blinded observer used ImageJ to trace and measure cluster area. Results indicate the average area of at least 100 unique clusters per treatment group. To quantify cleaved-caspase 3 positivity, a blinded observer analyzed cross sections of fixed, sectioned, and stained PDAC clusters. The number of nuclei as well as positive and negative staining was recorded in at least 50 cross sections of unique PDAC cell clusters from 3 biological replicate experiments.

### TUNEL Staining and Quantification

The sections from two biological replicate experiments were stained using a Terminal deoxynucleotidyl transferase (TdT) dUTP Nick-End Labeling (TUNEL) assay (Millipore S7165), according to the manufacturer’s instructions. The slides were deparaffinized, pretreated, and antigen retrieval was done by first using a TdT enzyme and then a rhodamine antibody solution. DAPI was used as a nuclear counterstain. Images of the fluorescently labelled sections were obtained using a Leica SP5X Upright Two-Photon Confocal Microscope. To quantify TUNEL positivity, the number of TUNEL positive nuclei were obtained as well as total number of nuclei per cell cluster.

## Acknowledgments

We thank Carlos Espinoza and Daniel Long for technical supports. We thank all the members of the PanTErA (Pancreatic Tumor Eradication Association) at the University of Michigan for constructive suggestions and feedback. We thank Drs Benjamin L Allen, Elizabeth Lawlor and Ron Koenig for serving of Dr Brannon’s thesis committee and for sharing their expertise and input on this project.

