## Supplementary figures and images for "Beta 1 Integrin Signaling Mediates Pancreatic Ductal Adenocarcinoma Resistance to MEK Inhibition"

### Supplemental Figure 1

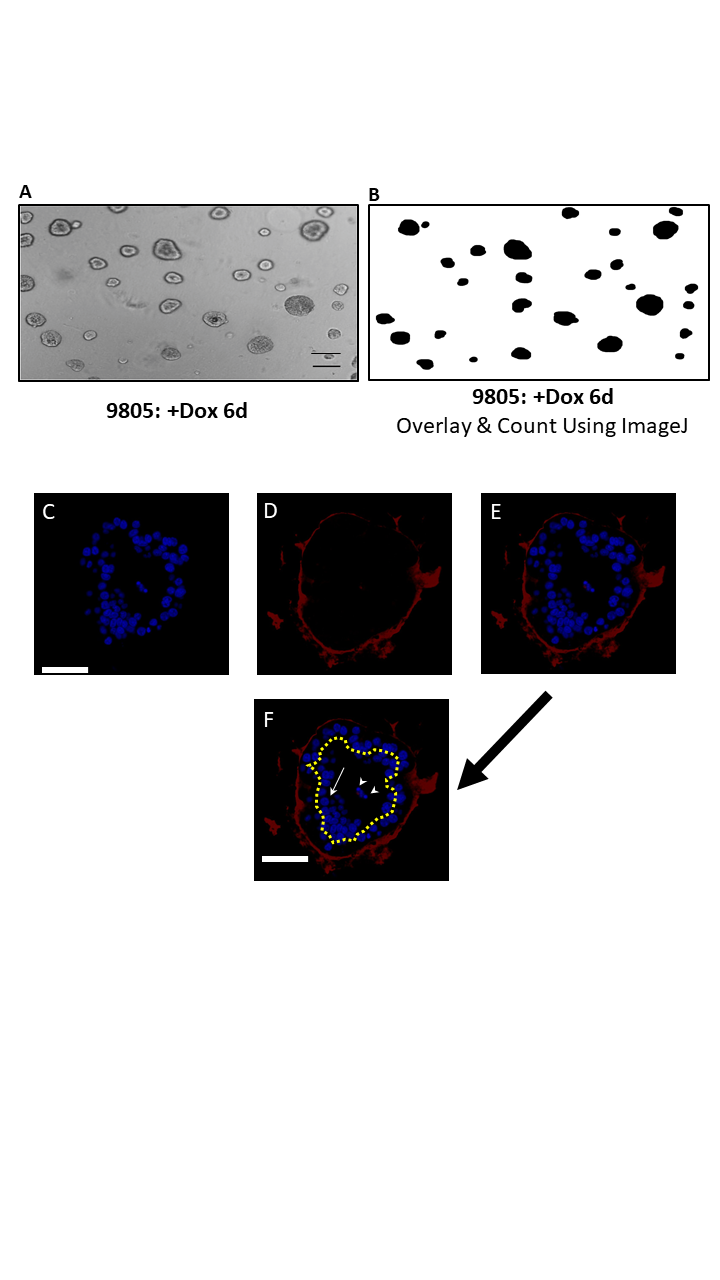

### Supplemental Figure 2

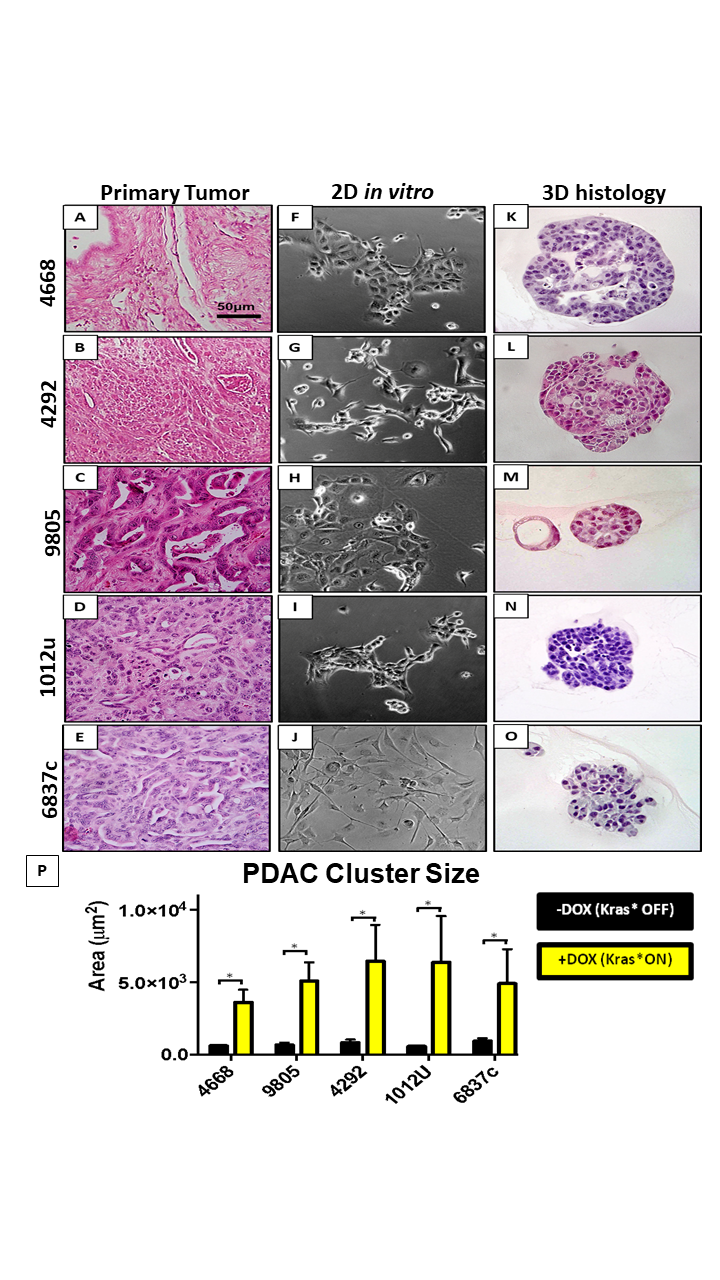

### Supplemental Figure 3

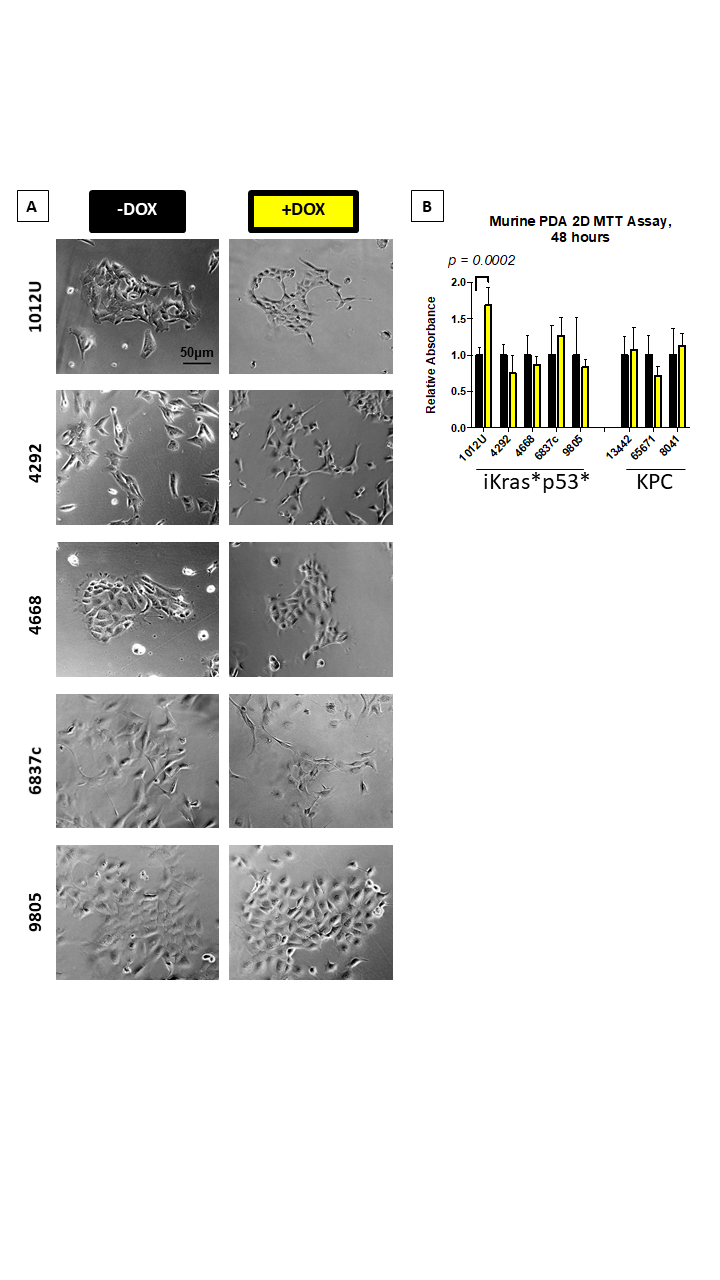

### Supplemental Figure 4

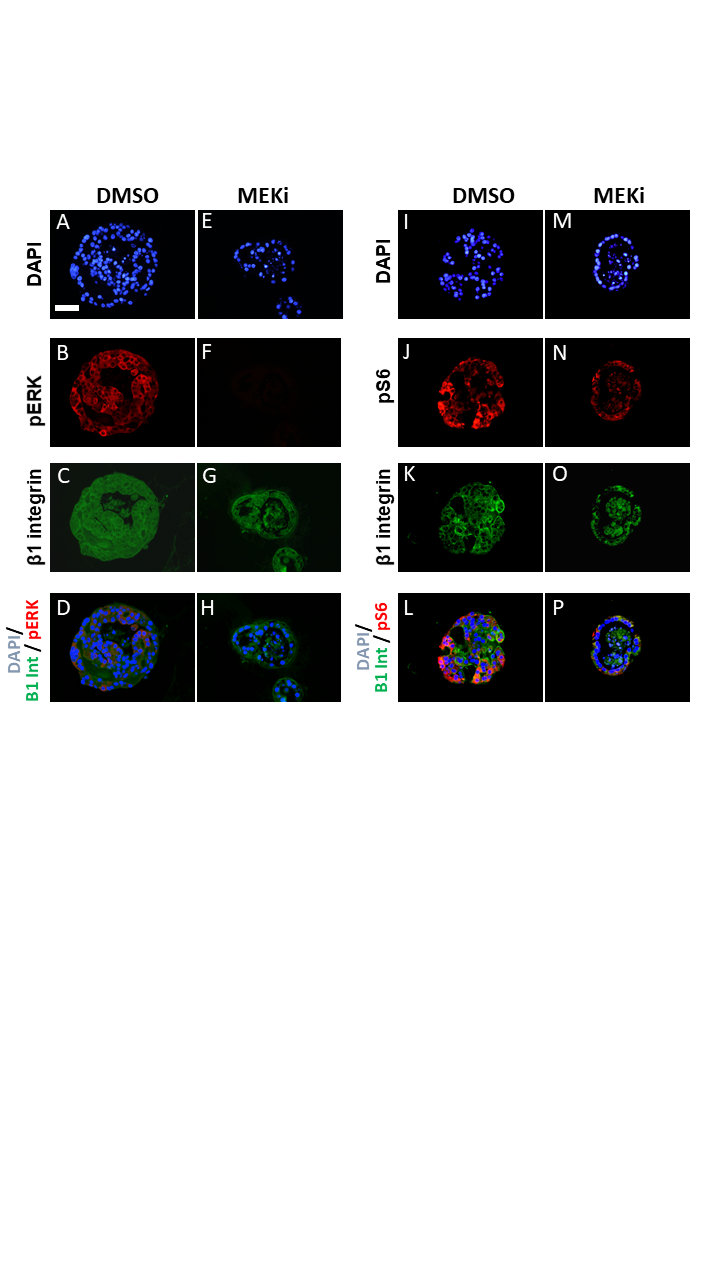

### Supplemental Figure 5

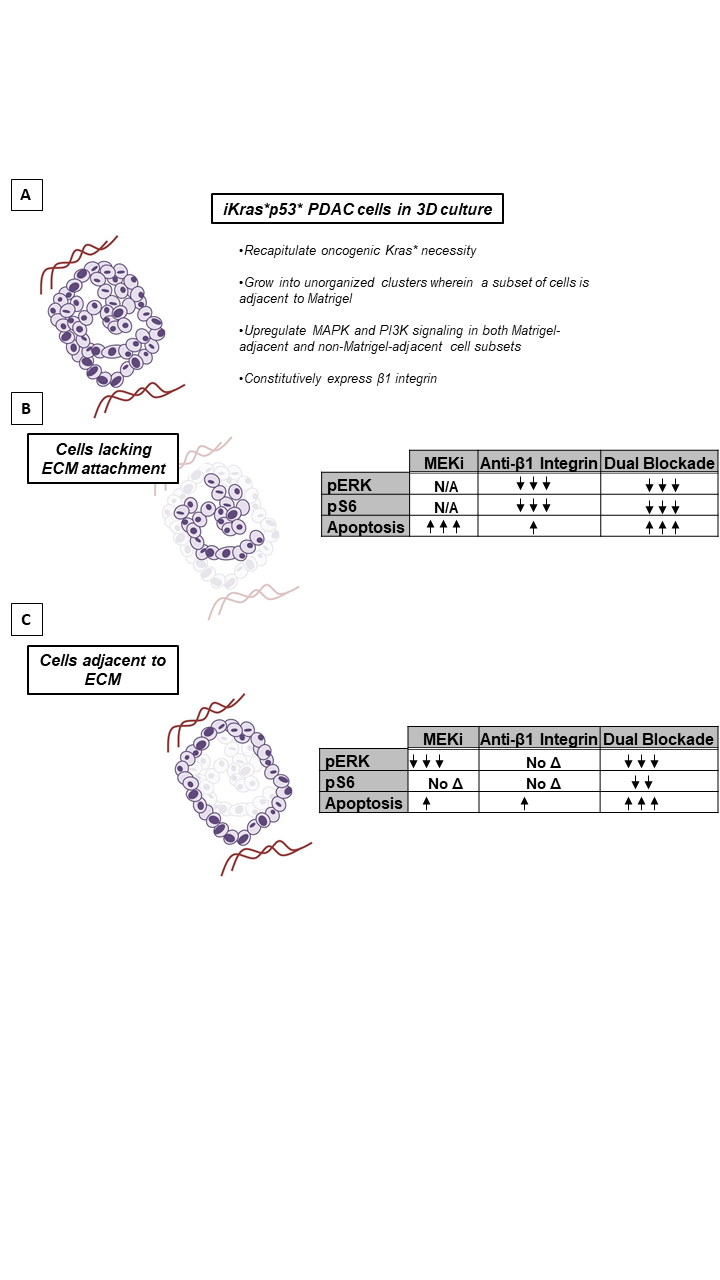

### Supplemental Figure 6

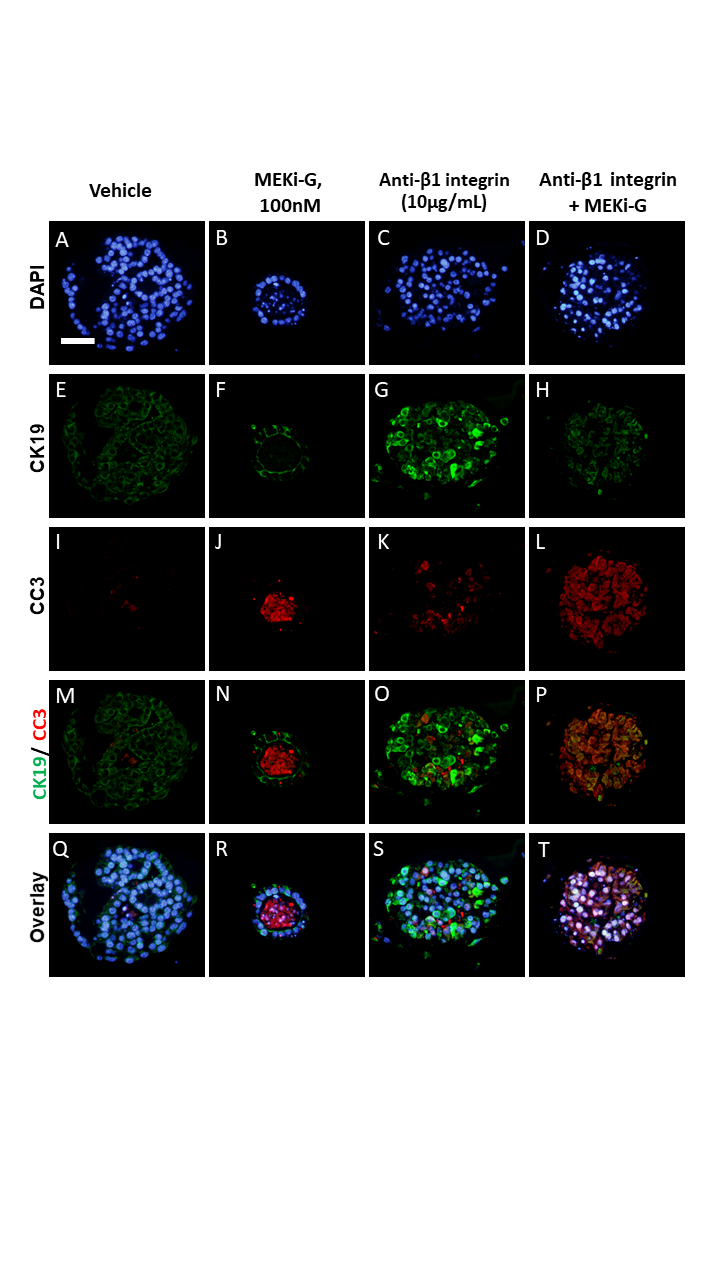

### Supplemental Table 1

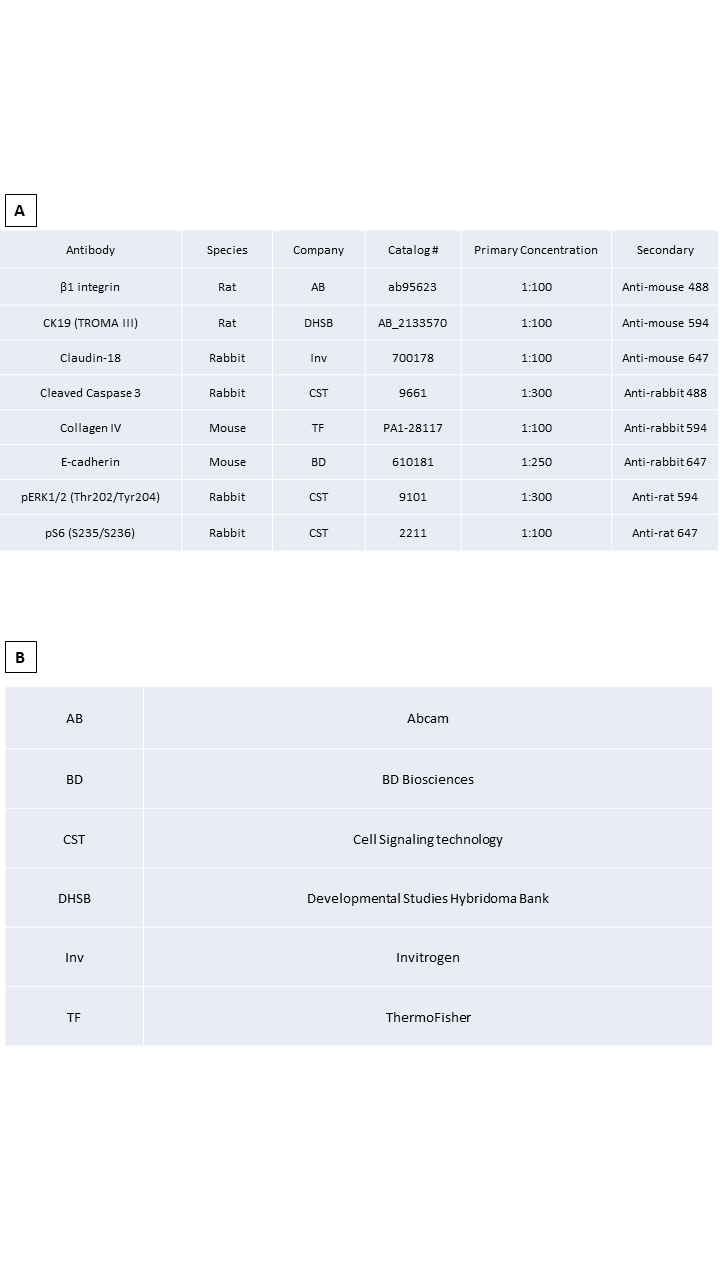
