## Supplemental Figure Legends for "Beta 1 Integrin Signaling Mediates Pancreatic Ductal Adenocarcinoma Resistance to MEK Inhibition"

**Supplemental Fig. 1** Quantifying 3D clusters and Matrigel-adjacent cells. **a-b** Measuring cluster area as an indication for size. Scale bar: 250 µm **c-e** Nuclei were quantified using type IV collagen-stained slides. Matrigel-adjacent nuclei were quantified separately. Scale bar: 50µm **f** Cluster cross sections were defined as groupings of >4 nuclei with visible nucleoli.

**Supplemental Fig. 2** PDAC cells harvested from in vitro assays display characteristics of primary tumor histology. **a-e** Hematoxylin/eosin stain of primary iKras*p53* PDAC tumors. **f-j** Brightfield images of PDAC cell lines **k-o** Hematoxylin/eosin stain of iKras*p53* PDAC clusters. Scale bar: 50µm. **p** Quantification of cluster area size. *$p<0.01$ in Student's t test analysis.

**Supplemental Fig. 3** iKras*p53* PDAC cell lines fail to recapitulate oncogenic Kras* dependency in 2D. **a** Brightfield images of cells with Kras* off or on, respectively. **b** Quantification of absorbance following MTT dye assay. Black bars represent Kras* off; yellow bars represent Kras* on. "P-value" is representative of Student's t-test result.

**Supplemental Fig. 4** MEK Inhibition does not affect β1 Integrin expression. **a, e, i, m** Nuclear visualization with DAPI. Scale bar: 50µm **b, f** Staining for phosphorylated ERK (pERK) **j, n** Staining for phosphorylated ribosomal s6 protein. **c, g, k, o** Staining for β1 integrin. **d, h, l, p** Overlay of all channels.

**Supplemental Fig. 5** β1 integrin signaling mediates oncogenic Kras* effector pathway signaling and resistance to MEK inhibition. **a** PDAC clusters in a 3D system recapitulate important aspects of PDAC. **b** Cells lacking ECM attachment are sensitive to MAPK inhibition. **c** Cells adjacent to the ECM displayed a survival advantage when MEK is inhibited.

**Supplemental Fig. 6** β1 integrin blockade sensitizes PDA cells to MEKi-induced apoptosis. **a-d** Nuclear visualization with DAPI. Scale bar: 50µm **e-h** CK19 staining indicating a ductal phenotype. **i-l** Cleaved caspase 3 (CC3) indicating apoptosis. **m-p** Merged channel images of CK19 and CC3 staining. **q-t** Overlay of DAPI, red, and green channels.

**Supplemental Table 1** Tables containing information regarding the antibodies used for immunofluorescent staining. **a** Contains information regarding the type of primary antibody used at the specified concentrations and the type of secondary antibody used. **b** Table of abbreviations used for companies supplying antibodies.
